# High-yield expression and purification of the non-phosphorylated form of Human CaMKIIα Kinase

**DOI:** 10.64898/2026.10.02.756153

**Authors:** Shubham Yadav, Latika Dina Gaude, Santosh Kumar

**Author notes:** Author for correspondence, (S.K.).

## Abstract

Human Ca^2+^/calmodulin (CaM)-dependent protein kinase II alpha (CaMKIIα) is a highly abundant and conserved protein in the brain. It is a multifunctional serine/threonine kinase that phosphorylates cellular substrates across subcellular compartments, activating signaling pathways essential for long-term memory storage. However, a detailed understanding of protein structure and enzyme-substrate interaction mechanisms require a method to produce active, stable, and pure proteins at scale. Since phosphorylation can alter catalytic activity and substrate affinity, examining the non-phosphorylated state enables an unbiased characterization of basal enzymatic function. Therefore, producing the non-phosphorylated form of the kinase is important for biochemical and structural studies aimed at understanding kinase activation mechanisms.

We describe a detailed protocol for the expression and high-yield purification of full-length, non-phosphorylated human CaMKIIα in bacterial cells. CaMKIIα was co-expressed with λ-phosphatase to maintain it in a non-phosphorylated state. Protein expression was induced by autoinduction at low temperature to enhance the stability and solubility of CaMKIIα. Recombinant CaMKIIα was purified by Ni-NTA affinity chromatography, followed by gel filtration chromatography. Although CaMKIIα has been considered difficult to purify in its non-phosphorylated form, we obtained it in high yield, and the protein was predominantly non-phosphorylated. Our results show that combining co-expression with autoinduction improves both the solubility and yield of the homogeneous non-phosphorylated form of CaMKIIα, which could be suitable for subsequent biochemical and structural studies.

**Highlights:**

- Human CaMKIIα was successfully expressed in *E. coli* using autoinduction.
- Co-expression with λ-phosphatase maintained CaMKIIα predominantly in a non-phosphorylated state.
- The purified CaMKIIα showed no detectable Thr286 phosphorylation and retained Ca^2+^/calmodulin-dependent kinase activity.

## Introduction

Calcium/calmodulin-dependent protein kinase II (CaMKII) is a highly conserved Ser/Thr kinase that plays a pivotal role in neuronal signaling and synaptic plasticity **(Griffith, 2004; Colbran et al., 2004; Yasuda et al., 2022)**. In vertebrates, CaMKII is encoded by four genes (α, β, γ, and δ), which produce >25 distinct isoforms through alternative splicing. The α and β isoforms mainly in the brain form large holoenzymes, typically a mix of α and β subunits, assembling into dodecameric (12) or sometimes tetradecameric (14) structures **(Hoelz et al., 2003; Rosenberg et al., 2006; Rellos et al., 2010)**. Each CaMKII subunit comprises an N-terminal catalytic kinase domain, an autoinhibitory regulatory (pseudo-substrate) segment, a variable linker, and a C-terminal association (hub) domain that mediates oligomerization **(Nguyen et al., 2015)**. Under resting conditions, the autoinhibitory segment blocks the kinase active site; binding of Ca^2+^/calmodulin relieves this inhibition and activates each subunit, thereby exposing the critical Thr286 (α-isoform numbering) residue in the regulatory domain. Once Thr286 is exposed, it can be autophosphorylated by a neighboring subunit, trapping the enzyme in a Ca^2+^-independent active state **(Bhattacharyya et al., 2020)**. This molecular switch mechanism underlies CaMKII’s role in long-term potentiation (LTP) and memory **(Lisman et al., 2012; Tao et al., 2021)**.

Purifying functional CaMKII kinase often poses general challenges, including low yield, instability, and heterogeneity due to its large, multi-domain structure, which increases susceptibility to proteolysis and aggregation **(Bolton et al., 2024)**. Expressing CaMKII in bacterial cells often yields fragments and aggregates unless conditions are optimized **(Waxham et al., 1989; Praseeda et al., 2004)**. Additionally, CaMKII autophosphorylation at Thr286, along with inhibitory sites Thr305/306, readily occurs during expression and purification, generating a heterogeneous mixture of activation states (**Bhattacharyya et al., 2020; Lučić et al., 2023)**. Many biochemical and structural studies rely on the autoinhibited (non-phosphorylated) form, contaminating phospho-CaMKII can exhibit aberrant activity, complicating interpretation and posing a major obstacle **(Lučić et al., 2023)**. In fact, short incubations of CaMKII in extract or on CaM-affinity columns can lead to inadvertent phosphorylation unless phosphatases are present **(Bolton et al., 2024)**. Consequently, isolating homogeneous, non-phosphorylated CaMKII requires extreme care.

Previous research has explored various methods for producing full-length CaMKIIα using bacterial, insect, and mammalian systems. Using insect cells, Brickey et al. obtained 12-15 mg/L via ammonium sulfate precipitation and calmodulin-Sepharose chromatography **(Brickey et al., 1990)**, while Török et al. achieved ∼11 mg/L via calmodulin affinity and anion-exchange chromatography **(Török et al., 2001)**. Other insect-cell approaches incorporated additional steps, including sucrose density centrifugation, cation- or anion-exchange chromatography, and gel filtration, to improve purity and enrichment of the holoenzyme **(Putkey et al., 1996; Kolb et al., 1998; Kolodziej et al., 2000; Rosenberg et al., 2005; Bolton et al., 2024)**. These studies show that insect cells effectively produce recombinant CaMKIIα; however, the process is labor-intensive, costly, less scalable, variable, and complex, with undefined phosphorylation states that complicate analyses of activity and structure.

Bacterial expression with IPTG induction using affinity-tagged constructs has been explored. Early bacterial studies showed that CaMKIIα could be produced in *E. coli;* however, quantitative yields of the heterogeneous phosphorylation of CaMKIIα were not reported **(Waxham et al., 1989)**. Moreover, Chao et al. reported the expression of recombinant CaMKIIα, followed by multiple chromatography steps; however, the yield was only 2.5 μg/L **(Chao et al., 2010)**. Recently, Bolton et al. optimized bacterial expression and purification using cation-exchange and calmodulin-affinity chromatography, reporting a substantially improved yield of approximately 4 mg/L (**Bolton et al., 2024)**. However, the preparation has notable limitations. The CaM-affinity step mainly enriches Ca^2+^/calmodulin-binding species, creating a bias towards partially activated forms and not fully resolving phosphorylation heterogeneity, leaving the enzyme’s phosphorylation state uncertain. Importantly, most purification strategies priorities recovery and purity rather than actively preventing phosphorylation of CaMKIIα during heterologous expression.

Autoinduction using lactose-based systems has emerged as a preferred alternative to IPTG induction, particularly for producing difficult recombinant proteins that often misfold, aggregate, or show toxicity in *E. coli* **(Studier 2005; Tahara et al., 2021; Fuente-Kratzborn et al., 2025)**. The autoinduction method has proven effective on various challenging targets, including amyloid precursor protein C99 **(Sharma et al., 2025)**, consensus interferon-α **(EL-Baky et al., 2015)** cysteine-rich engineered proteins such as protein G-5xCys **(Wang et al., 2021)**, and enzymes and receptor domains such as Trx1, GSTA1, MsrB, and T1R1 **(Briand et al., 2016)**.

Our approach builds on previous recombinant expression strategies while addressing a key limitation of CaMKIIα production, i.e., phosphorylation heterogeneity during protein expression. We co-expressed human CaMKIIα with λ phosphatase in *E. coli* using an autoinduction-based expression system. Concurrent expression of λ-phosphatase maintains CaMKIIα predominantly in its non-phosphorylated, autoinhibited state. The resulting His-tagged CaMKIIα holoenzyme was purified using a two-step procedure comprising Ni^2+^-affinity chromatography followed by gel filtration chromatography, yielding a highly pure, full-length holoenzyme. The purified protein was free of detectable Thr286 phosphorylation, making it suitable for further biochemical and structural studies, and it retained catalytic activity towards substrates, indicating that the protein was properly folded and functionally competent.

## Result and Discussion

### Expression, Purification and Biochemical Validation of Recombinant His-Tagged CaMKIIα

To obtain purified recombinant CaMKIIα, His-tagged CaMKIIα was expressed in bacterial cells using an autoinduction method followed by purification through Ni-NTA affinity and gel-filtration chromatography (GFC) **(Fig. 1A)**. SDS-PAGE analysis of whole-cell lysates showed efficient expression of recombinant CaMKIIα after autoinduction, with a prominent 55-60 kDa band in induced lysate **(Fig. 1B, lane L1)**, whereas this band was less visible in pre-induced lysate **(Fig. 1B, lane L2)**. Lysate was subjected to Ni-NTA affinity chromatography to enrich His-tagged proteins, yielding ∼5 mg/L. The enriched proteins were further purified by GFC, which resolved two peaks, P1 and P2, with P1 containing the major protein and P2 corresponding to a smaller protein population that elutes at a higher retention volume **(Fig. 1C)**. Fractions from P1 were pooled, yielding ∼3 mg/L of total protein. SDS-PAGE analysis showed substantial enrichment of a protein at the expected CaMKIIα size of 55–60 kDa, with minor impurities **(Fig. 1D**), indicating increased purity. Immunoblotting with anti-His confirmed the protein’s identity, showing a single band at 55– 60 kDa, consistent with full-length recombinant CaMKIIα **(Fig. 1E)**. Consistent with these findings, immunoblotting with a CaMKIIα-specific antibody detected a single major band of identical electrophoretic mobility in the purified sample **(Fig. 1F)**, confirming that the purified protein was recombinant CaMKIIα.

**Figure 1:**
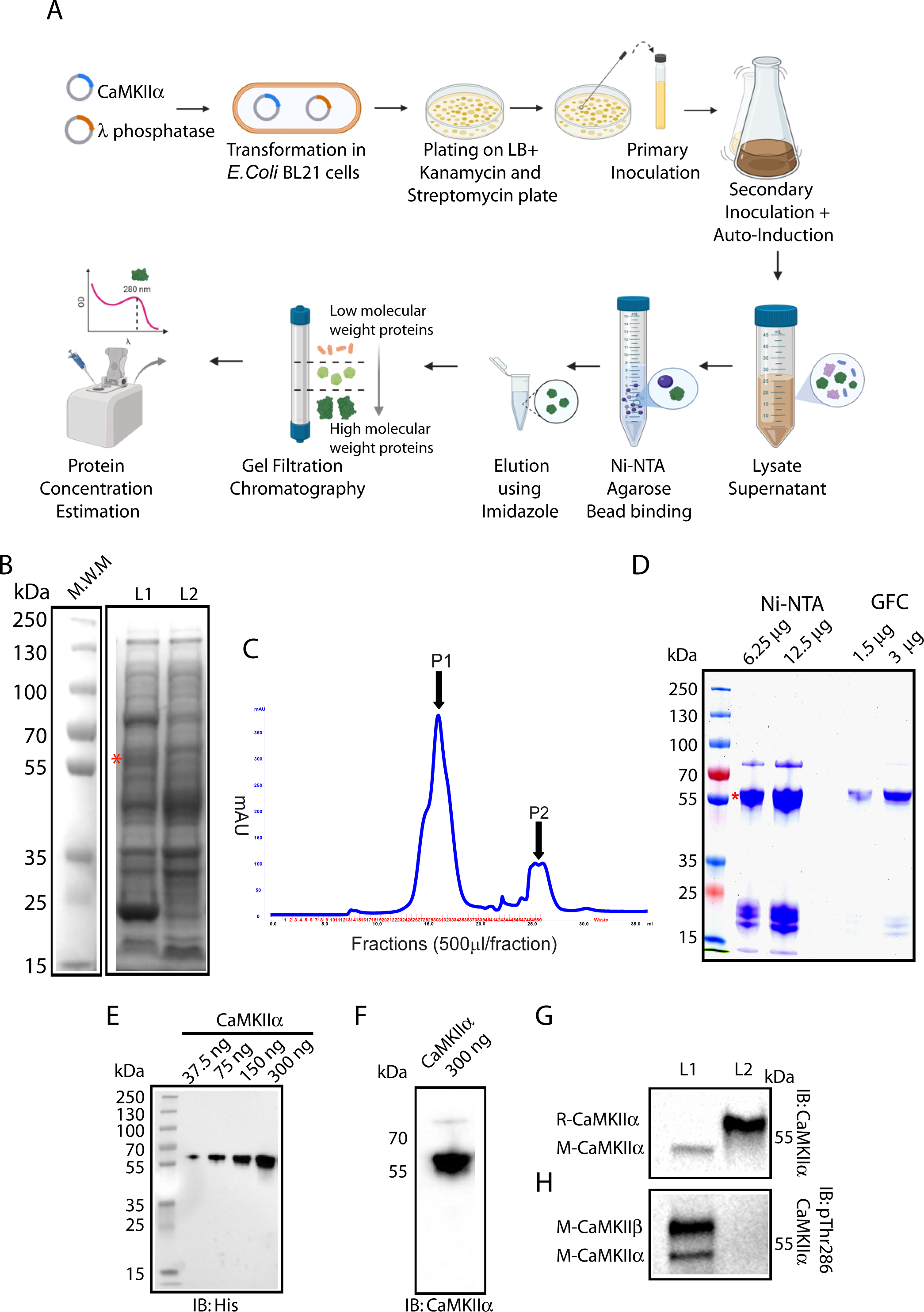
Expression and purification of human CaMKIIα protein in bacterial cells. A: A schematic illustrates the expression and purification of human CaMKIIα in bacterial cells. B: Protein expression was analyzed by loading cell lysates equivalent to OD_600_ = 0.1/lane and separating them on a 10% SDS-PAGE gel. L1 shows the autoinduced cells, whereas L2 corresponds to non-induced control cells. The red asterisk indicates the expressed CaMKIIα protein. C-D: After autoinduction, cells were lysed, and CaMKIIα-6-His was purified by Ni-NTA affinity and gel filtration chromatography. The 500μl fraction/tube from peak P1 (fractions 26-35) was collected, pooled (C), and both Ni-NTA- and gel-filtration column-purified CaMKIIα-6-His were analyzed by SDS-PAGE (D). E-F: To verify the CaMKIIα-6-His purified fraction, protein samples were separated by SDS-PAGE, and the bands were transferred to a nitrocellulose membrane and probed with mouse monoclonal anti-His (E) or anti-CaMKIIα (F) antibodies and detected the ∼55-60 kDa molecular-weight band of CaMKIIα. G: To determine the phosphorylation state of purified recombinant CaMKIIα, 600 ng of purified CaMKIIα was separated by SDS-PAGE gel (L2) and then subjected to a western blot, probed with mouse monoclonal CaMKIIα, which recognizes both phosphorylated and non-phosphorylated forms of CaMKIIα. As a control, 10 μg of mouse brain lysate (MBL) was run in parallel (L1). H: In panel G, blot was reprobed using a Rabbit monoclonal Thr286P-CaMKII antibody, which recognizes phosphorylation on both CaMKIIα and CaMKIIβ. Phosphorylation signals were detected in mouse CaMKIIα and CaMKIIβ isoforms, but the band for purified recombinant CaMKIIα was not present. Bands for total mouse CaMKII and recombinant CaMKIIα were both visible, with’ M’ indicating mouse and’ R’ indicating recombinant.

Previously, CaMKIIα was produced in bacterial systems using IPTG induction, resulting in relatively low protein yields **(Chao et al., 2010; Bhattacharyya et al., 2016)**. This was probably due to suboptimal induction timing and metabolic stress on host cells **(Donovan et al., 1996)**, leading to protein misfolding and aggregation, with the aggregates remaining in the insoluble fraction **(Baneyx and Mujacic, 2004; Studier, 2005; Sørensen and Mortensen, 2005)**. Consistent with these findings, this limitation hindered a straightforward comparison between IPTG induction and autoinduction in our study. In contrast, we employed the autoinduction method, which produced much higher CaMKIIα (∼3 mg/L) and more consistent expression. Importantly, N-terminal SUMO tagging of CaMKIIα enhanced solubility without altering its structural integrity or oligomeric state **(Bhattacharyya et al., 2016)**. Earlier reports have demonstrated that kinases and other multi-domain or complex proteins are more efficiently expressed and purified using autoinduction methods than with traditional IPTG induction **(Studier, 2005; Terpe, 2006; Rosano and Ceccarelli, 2014)**. For instance, the ASK1 kinase domain has been successfully expressed in a soluble form using autoinduction. In contrast, IPTG induction often yields low amounts or inactive proteins **(Volynets et al., 2016)**.

To test the phosphorylation status of CaMKIIα, we used Western blot analysis with a phospho-specific CaMKII antibody and an antibody that detects total CaMKIIα protein. No signal was observed with the phospho-specific antibody in recombinant CaMKIIα, indicating that phosphorylated CaMKIIα protein was not present **(Fig. 1G, lane L2)**. Mouse brain lysate was used as a control, confirming the detection of both brain-specific phosphorylated isoforms, CaMKIIα and CaMKIIβ, by phospho-specific CaMKII antibody **(Fig 1G, lane L1)**. Using the CaMKIIα antibody, which detects both phosphorylated and non-phosphorylated CaMKIIα, bands corresponding to mouse CaMKIIα and recombinant non-phosphorylated CaMKIIα were observed, confirming CaMKIIα expression **(Fig. 1H)**. Taken together, these results demonstrate successful expression, purification, validation, and enrichment of non-phosphorylated recombinant CaMKIIα, enabling structural, functional, and in vitro phosphorylation studies.

Most in vitro assays include Ca^2+^/CaM to stabilise CaMKIIα, inadvertently triggering autophosphorylation at Thr286 and converting the protein to its phosphorylated form **(Hashimoto et al., 1987; Giese et al., 1998)**. Consequently, maintaining CaMKIIα in a non-phosphorylated form during purification and analysis is a major challenge, as basal or trace Ca^2+^/CaM can induce autophosphorylation (**Bolton et al., 2024)**. This explains why previous studies have failed to convincingly isolate stable, non-phosphorylated CaMKIIα **(Putkey et al., 1996; Kolb et al., 1998; Kolodziej et al., 2000; Rosenberg et al., 2005)**.

Obtaining stable, non-phosphorylated CaMKIIα provides advantages for studying its initial activation, CaM binding, autophosphorylation, inhibitory phosphorylation, and basal conformational states without phosphorylation artefacts, improving understanding of its regulation and role in cellular signaling. Our autoinduction strategy with λ phosphatase inhibits autophosphorylation at Thr286, maintaining CaMKIIα in a non-phosphorylated, autoinhibited state. A similar method with IPTG induction has been used for purification, but yields are low **(Chao et al., 2010)**. Recently, Bolton et al. showed the presence of non-phosphorylated CaMKIIα in cell lysates. The authors reported high yields using CaM-based affinity chromatography **(Bolton et al., 2024)**, but trace Ca^2+^ likely caused autophosphorylation during purification, with the phosphorylation status unreported. Our approach allows large-scale purification of non-phosphorylated CaMKIIα via autoinduction, avoiding CaM affinity steps and calcium exposure, thus preserving its native state. Thus, our study offers a major advantage for studying the intrinsic biochemical properties, activation dynamics, and regulatory mechanisms of CaMKIIα without interference from prior autophosphorylation.

### Ca^2+^/Calmodulin-dependent Activation of CaMKIIα

Having established the successful purification of recombinant non-phosphorylated CaMKIIα **(Fig. 1)**, we next examined whether the purified kinase retained its catalytic activity. CaMKIIα exists in an autoinhibited conformation in the absence of Ca^2+^/calmodulin (CaM). Binding of the Ca^2+^/CaM complex relieves this inhibition, enabling intermolecular autophosphorylation at Thr286, a hallmark of kinase activation that confers autonomous catalytic activity **(Fig. 2A)**. To facilitate this activation, recombinant calmodulin was purified separately and analyzed by SDS-PAGE, revealing a single predominant protein band migrating at approximately 17 kDa, consistent with the expected molecular weight of calmodulin and indicative of a purified preparation suitable for kinase activation assays **(Fig. 2B)**. Incubating non-phosphorylated CaMKIIα with Ca^2+^ and CaM showed strong phospho-Thr286 signals **(Fig 2C, lane L2)** compared to control **(Fig 2C, lane L1)**. Total CaMKIIα levels remained consistent **(Fig. 2D)**, confirming phosphorylation was due to activation. Thr286 is the principal autophosphorylation site that stabilizes the active conformation of CaMKIIα after Ca^2+^/CaM binding and is widely accepted as a biochemical marker of enzymatically active CaMKIIα **(Miller and Kennedy, 1986; Hudmon and Schulman, 2002)**. Therefore, the presence of phospho-Thr286 with constant total CaMKIIα indicates that the purification preserved the kinase’s structural and catalytic integrity, suitable for substrate phosphorylation, interaction, and structural studies. These results support autophosphorylation as a reliable indicator of CaMKIIα’s physiological function and stability **(Kim et al., 2001; Hudmon and Schulman, 2002; Lisman et al., 2012; Coultrap and Bayer, 2012)**. Taken together, our results confirm that the purified, homogeneous, non-phosphorylated form of CaMKIIα is functionally active, making it suitable for further biochemical and structural research, as well as *in vitro* phosphorylation assays to identify and characterize its downstream substrates.

**Figure 2:**
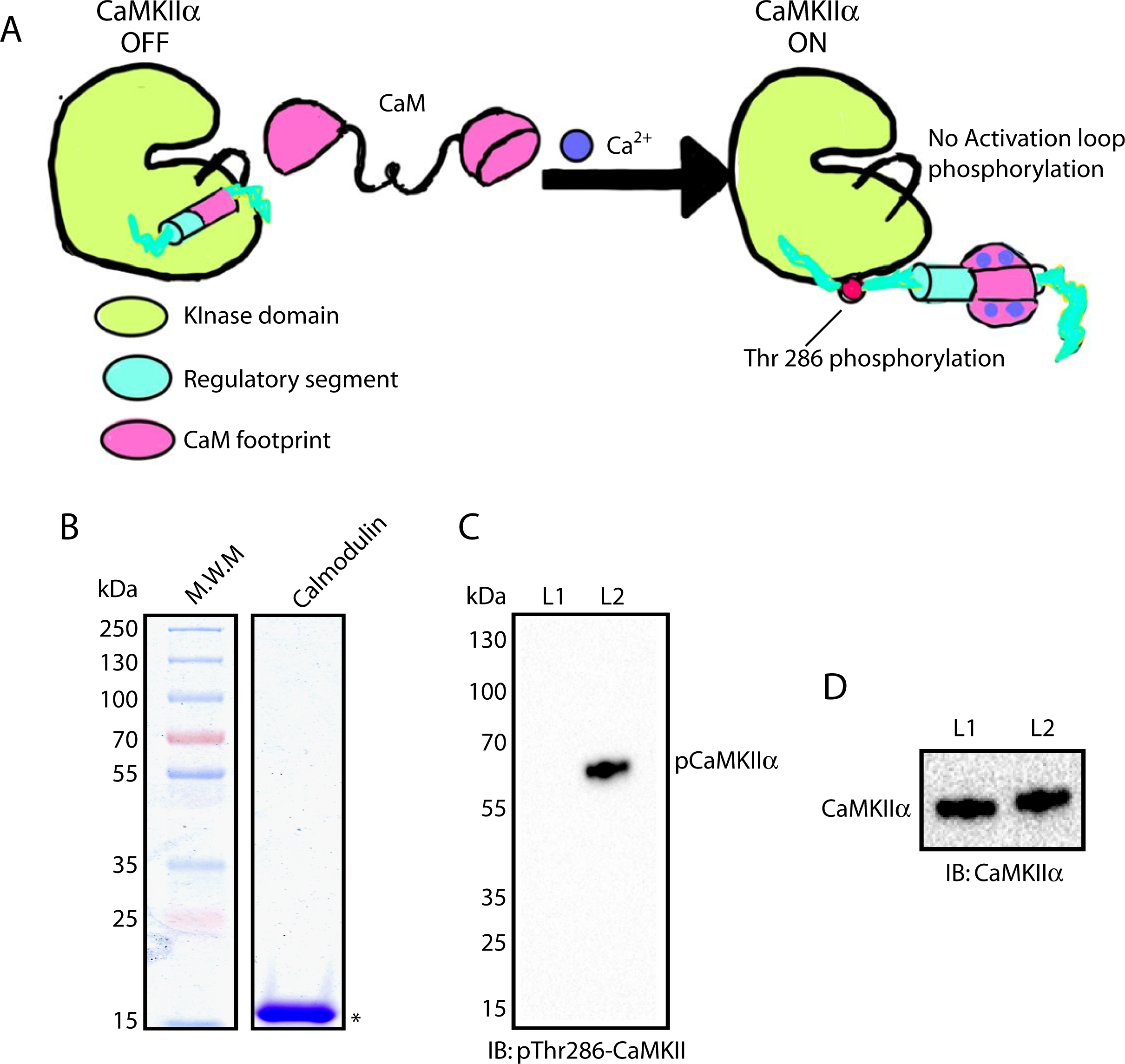
Purified non-phosphorylated form of CaMKIIα is functionally active. A: Schematic illustrates the phosphorylation of CaMKIIα at threonine (Thr) 286, mediated through CaM and Ca^2+^. B: CaM was expressed in bacterial cells using the autoinduction method and purified using glutathione agarose beads. 7 μg of purified CaM was analysed on a 10% SDS-PAGE gel. C-D: The purified non-phosphorylated CaMKIIα was incubated with a reaction mixture containing 5 μM CaM, 100 μM ATP, and 1 mM CaCl_2_ in 100 μl of buffer to convert it into phosphorylated CaMKIIα (Thr286P) (lane L2). As a control, the non-phosphorylated CaMKIIα was incubated with the same buffer without added reagents (lane L1). Phosphorylation was measured by Western blotting using a rabbit monoclonal Thr286P-CaMKII antibody (C), and total CaMKIIα was detected with a mouse monoclonal CaMKIIα antibody (D). 10 μl samples were loaded for Western blot analysis in lanes L1 and L2.

## Materials And Methods

### Reagents, Plasmids and bacterial strains

All reagents, chemicals and other information are listed in Key Resources Supplemental **Table S1**. pSTM3-CaMKIIα, pCDFDuet1-pptase and pCaM vectors **(Chao et al., 2011; Bhattacharyya et al., 2016)** were generously provided by Prof. John Kuriyan’s laboratory (Vanderbilt University, USA). In the pSTM3-CaMKIIα vector, the N and C-terminal domains of human full-length CaMKIIα were tagged with Sumo and 6-histidine, respectively **(Bhattacharyya et al., 2016)**. pCDFDuet1-pptase vector expresses Lambda Phosphatase. CaM (Calmodulin) was amplified from the pCaM vector using primers listed in Table S1 and subcloned into the pMD49 vector containing an N-terminal GST-6xHis-TEV expression tag. All the experiments were done twice unless otherwise specified. We used mouse brain lysate as a control in one experiment. It was prepared previously and also used in this study. All the details are mentioned in our previous study **(Yadav et al., 2025)**.

### Expression and purification of CaMKIIα and Calmodulin

We co-expressed CaMKIIα with Lambda Phosphatase in *E. coli* BL21 (DE3) cells using the autoinduction method **(Seeliger et al., 2005; Chao et al., 2010)**. Briefly, the expression construct was co-transformed into *E. coli* BL21 cells and cultured overnight at 37°C with shaking at ∼260 rpm in 100 ml of Luria–Bertani (LB) medium supplemented with Kanamycin at 50 µg/ml and Streptomycin at 100 µg/ml. The next day, an aliquot corresponding to an OD_600_ of 1.0 was collected as the pre-induction sample, and the remaining culture was inoculated into 1 litre of Terrific Broth (TB) media (for g/l: tryptone 12 g, yeast extract 24 g, 0.4% glycerol, 9.4 g Potassium phosphate dibasic, and 2.2 g Potassium phosphate monobasic) supplemented with Kanamycin at 50 µg/ml and Streptomycin at 100 µg/ml, followed by incubation at 37°C for 1 hour. Protein expression was induced with 0.2% lactose, and the culture was grown at 37°C for an additional 4 hours, then shifted to 16°C for 18 hours. Cells were harvested the following day, and a sample with an OD_600_ of 1.0 was collected as the post-induction sample. Samples with an OD_600_ of 1.0 were pelleted and boiled in SDS-loading buffer. To check protein expression, a 0.1 OD was loaded into a well, and proteins were separated on a 10% SDS-PAGE gel.

For protein purification, cells were pelleted and resuspended in ice-cold 15 ml of lysis buffer (50 mM HEPES, pH 7.4, 150 mM NaCl, 10% glycerol, 1 mM DTT, 30 mM imidazole) containing 1 mM PMSF and the complete protease inhibitor cocktail (Roche #11873580001). Cells were lysed by passing them through a French press (Spectronic Instruments, model FA078) twice at 11000 PSI. Cell lysates were centrifuged at 100,000Xg for 40 minutes at 4°C and the clarified supernatant was transferred into new tubes. The protein concentration was measured using Bio-Rad Protein Assay Dye Reagent Concentrate (Bio-Rad, USA). Clarified supernatant was loaded on 1ml Ni-NTA column. Column was washed with washing buffer (50 mM HEPES pH 7.4, 300 mM NaCl, 10% glycerol, 50mM imidazole) and bound proteins were eluted using elution buffer (50 mM HEPES pH 7.2, 50 mM NaCl, 10% glycerol, 500mM imidazole). Protein-containing fractions were collected, pooled and concentrated to 550 μl and centrifuged at 100,000Xg for 10 minutes at 4°C before loading on the Gel filtration column Superose^®^ 6 10/300 GL (Cytiva, USA). Column was preequilibrated with running buffer (50 mM HEPES, pH 7.4, 100 mM NaCl, 10% glycerol). The flow rate was set to 500μl/min, and 500μl fractions were collected per tube. Protein-containing fractions from peak 1 (fractions 26 to 35) were pooled, and the concentration was measured using the Bio-Rad Protein Assay Dye Reagent, as described earlier.

Chicken calmodulin was also expressed in *E. coli* BL21 (DE3) cells using the autoinduction method, as previously. For calmodulin purification, cells were lysed in the lysis buffer described above, but without imidazole. Clarified supernatant was incubated with 1 mL glutathione agarose beads (Thermo Fisher Scientific #16100) on a rotamixer for 2h at 4°C. Beads were washed with washing buffer (50 mM HEPES pH 7.4, 150 mM NaCl, 10% glycerol), and proteins were eluted by overnight incubation with TEV-His protease (GenScript, USA) at 4°C on a rotamixer. Subsequently, eluted proteins were incubated with Ni-NTA-Agarose beads for 30 minutes at 4°C to remove TEV-His. The supernatant was collected, and protein concentration was measured using the Bio-Rad protein assay as before. Purified proteins were analyzed on a 10% SDS-PAGE gel.

### Gel electrophoresis and Western blot

All SDS-PAGE experiments used a 10% gel. Gels were stained with Coomassie Brilliant Blue R-250 dye. For the Western blot, protein samples were separated on a 10% SDS-PAGE gel, and bands were transferred to a 0.45m nitrocellulose membrane. Blots were probed with 1:1500-diluted mouse monoclonal anti-His (Invitrogen MA1-21315), 1:200-diluted mouse monoclonal anti-CaMKIIα (6G9) (Santa Cruz Biotechnology sc-32288), and 1:1000-diluted rabbit monoclonal anti-Thr286p-CaMKII (D21E4) (Cell Signaling Technology #12716) antibodies. Proteins were visualized using enhanced chemiluminescence with HRP–linked secondary antibodies at a dilution of 1:5000 for both anti-mouse (Bio-Rad, USA) and anti-rabbit (Bio-Rad, USA) antibodies, using the Super Signal West Pico PLUS Chemiluminescent Substrate (Thermo Scientific, USA).

### CaMKIIα phosphorylation

Non-Phosphorylated CaMKIIα was converted into Phosphorylated CaMKIIα (Thr286P) using a previously described method with some modifications **(Jama et al., 2009)**. Briefly, a reaction mixture containing 1 μM CaMKIIα, 5 μM CaM, 100 μM ATP, 1 mM CaCl_2_, was incubated for 10 min at 30 °C in the reaction buffer (50mM HEPES pH 7.4, 100mM NaCl, 10mM MgCl_2_). The reaction was performed in 100 μl. For the control, CaMKII was incubated with the reaction buffer. Western blot analysis was performed to detect total CaMKIIα and Phosphorylated CaMKIIα using mouse monoclonal CaMKIIα (Santa Cruz Biotechnology sc-32288) and Rabbit monoclonal Thr286P-CaMKII (Cell Signaling Technology #12716) antibodies, respectively.

## CRediT authorship contribution statement

**SY:** Investigation, Methodology, Writing; **LDG:** Investigation; **SK:** Supervision, Project administration, Conceptualization, Investigation, Methodology, Funding Acquisition, Writing and editing.

## Acknowledgments

We heartily thank all lab members for their continued and generous support and Nikhat Khan for skillful technical assistance. S.K.’s laboratory is funded by the Department of Biotechnology (DBT), Ministry of Science and Technology, India (BT/PR38584/MED/122/247/2020); the Department of Science and Technology (DST), Ministry of Science and Technology, India (CRG/2021/000732); and the DBT/Wellcome Trust India Alliance (IA/I/22/2/506480).

## Appendix A. Supplementary data

Table S1: Key Resources and Reagents used in the study

## Data availability

Data will be made available on request.

---

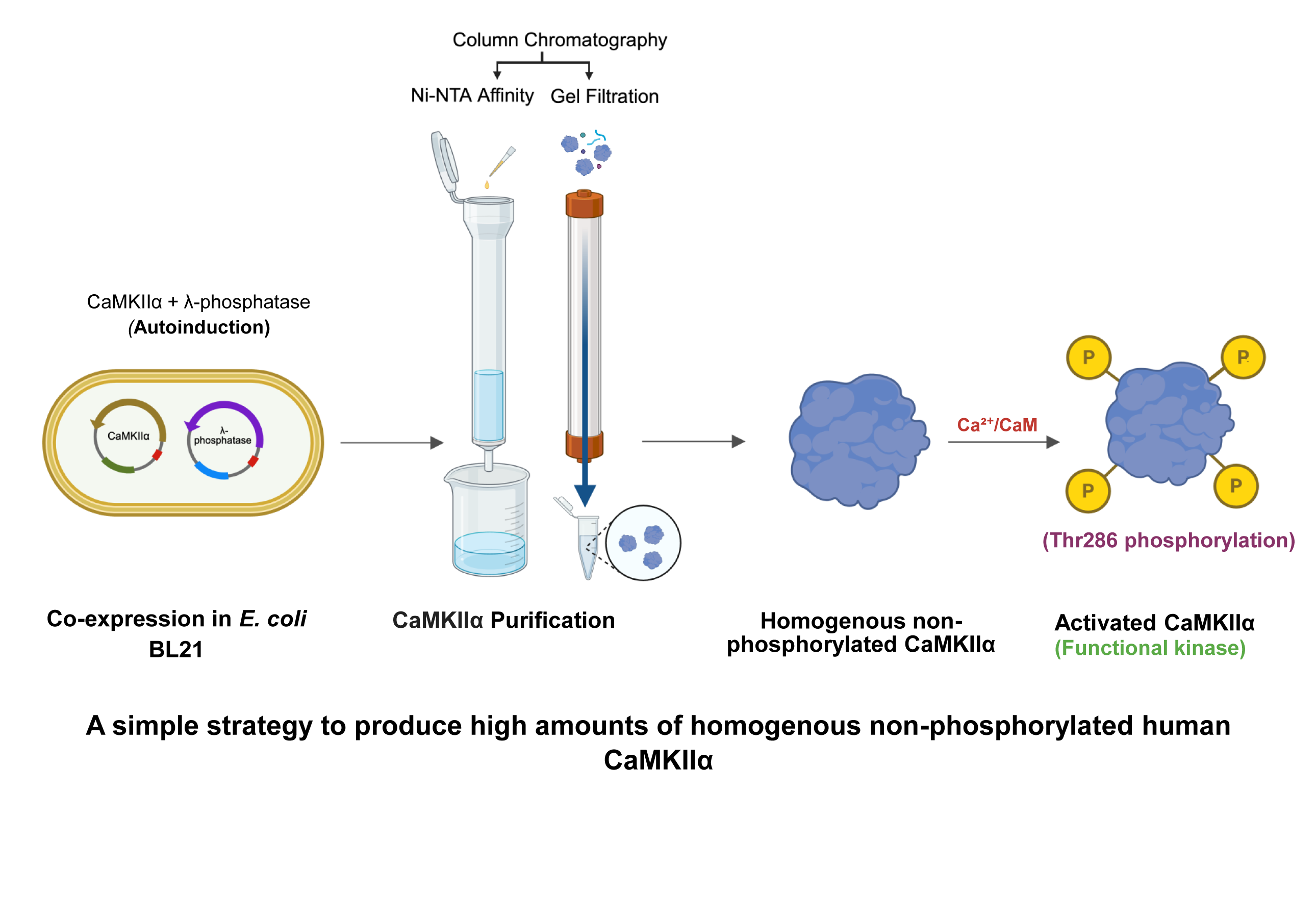

