## Supplementary material for "High-yield expression and purification of the non-phosphorylated form of Human CaMKIIα Kinase": Table S1

Table S1: Key Resources and Reagents used in the study

| Component | Source | Catalogue No. |
|---|---|---|
| **Antibodies** | | |
| Mouse monoclonal anti-His Antibody | Invitrogen | MA1-21315 |
| Mouse monoclonal anti-CaMKIIα Antibody (6G9) | Santa Cruz Biotechnology | sc-32288 |
| Rabbit monoclonal phospho-CaMKII (Thr286) Antibody (D21E4) | Cell Signalling Technology | 12716 |
| Goat anti-mouse HRP | Bio-Rad | 1706516 |
| Goat anti-rabbit HRP | Bio-Rad | 1706515 |
| **Chemicals** | | |
| Adenosine 5′-triphosphate disodium salt hydrate | Sigma-Aldrich | A26209 |
| Bio-rad Protein Assay Dye Reagent Concentrate | Bio-Rad | 5000006 |
| Calcium chloride dihydrate | Sigma-Aldrich | 223506 |
| Brilliant Blue R | Sigma-Aldrich | B0149 |
| Pierce™ Glutathione Agarose | Thermo Scientific | 16100 |
| Glycerol | Qualigens | Q15455 |
| HEPES (N-(2-hydroxyethyl)-1-piperazinee-N'-(2-ethanesulfonic acid) | Himedia | MB016 |
| Imidazole | Sigma-Aldrich | 56750 |
| D-Lactose Monohydrate | Sigma-Aldrich | 61339 |
| Ni-NTA Agarose | Qiagen | 30210 |
| PageRuler™ Plus Prestained Protein Ladder, 10 to 250 kDa | Thermo Scientific | 26619 |
| Phenylmethylsulfonyl Fluoride (PMSF) | US Biological Life Sciences | 329-98-6 |
| Potassium Phosphate Dibasic | Sigma-Aldrich | P8281 |
| Potassium Phosphate Monobasic | Sigma-Aldrich | 60229 |
| cOmplete™, EDTA free Protease Inhibitor Cocktail | Roche | 11873580001 |
| Sodium chloride | Qualigens | Q27605 |
| Superose® 6 10/300 GL | Cytiva | GE17-5172-01 |
| SuperSignal™ West Pico PLUS Chemiluminescent Substrate | Thermo Scientific | 34577 |
| TEV Protease, His | Genscript | Z03030 |
| Tryptone | HiMedia | CR014 |
| Yeast extract | HiMedia | RM027 |
| **Plasmids** | | |
| pSTM3-CaMKIIα | N and C-terminal domains of human full-length CaMKIIα tagged with Sumo and 6-histidine | Chao et al., 2011; Bhattacharyya et al., 2016 |
| pCDFDuet1-pptase | Expresses Lambda Phosphatase | Bhattacharyya et al., 2016 |
| pCaM | Template vector for CaM amplification | Bhattacharyya et al., 2016 |

| | | |
|---|---|---|
| pMD49-CaM | Contains GST-6×His-TEV-CaM | This study |
| **Primers** | | |
| CaM_F | CACGGTACCGAAAACCTGTATTTTCAGGGCATGGCTGACCAGCTGAC | This study |
| CaM_R | GATGCGGCCGCTCACTTTGCAGTCATCATCTGTAC | This study |
